# Molecular evolution of non-fertilizing sperm in Lepidoptera suggests minimal direct involvement in sperm competition

**DOI:** 10.1101/404236

**Authors:** Andrew J. Mongue, Megan E. Hansen, Liuqi Gu, Clyde E. Sorenson, James R. Walters

**Affiliations:** University of Kansas, Department of Ecology and Evolutionary Biology; North Carolina State University, Department of Entomology

**Keywords:** sperm dimorphism, population genetics, *Manduca sexta*, *Danaus plexippus*

## Abstract

Sperm are among the most variable cells in nature. Some of this variation results from non-adaptive errors in spermatogenesis, but many species consistently produce multiple sperm morphs, the adaptive significance of which remains unknown. Here, we investigate the evolution of dimorphic sperm in Lepidoptera, the butterflies and moths. Males of this order produce both fertilizing sperm and a secondary, non-fertilizing type that lacks DNA. Previous organismal studies suggested a role for non-fertilizing sperm in sperm competition, but this hypothesis has never been evaluated from a molecular framework. We combined published datasets with new sequencing in two species, the monandrous Carolina sphinx moth and the highly polyandrous monarch butterfly. Based on population genetic analyses, we see evidence for increased adaptive evolution in fertilizing sperm, but only in the polyandrous species. This signal comes primarily from a decrease in non-synonymous polymorphism in sperm proteins compared to the rest of the genome, suggesting stronger purifying selection, consistent with selection via sperm competition. Non-fertilizing sperm proteins, in contrast, do not show an effect of mating system and do not appear to evolve differently from the background genome in either species, arguing against the involvement of non-fertilizing sperm in direct sperm competition. Based on our results and previous work, we suggest that non-fertilizing sperm may be used to delay female remating in these insects and decrease the risk of sperm competition rather than directly affect its outcome.

## Introduction

Sperm cells display remarkable diversity throughout the animal kingdom (Pitnick, Hosken, & Birkhead, 2009), from small and plentiful to gigantic and few (Pizzari, 2006) to super-structure-forming (Higginson, Miller, Segraves, & Pitnick, 2012). This variation exists at every level, from fixed differences between species to variability within individual males (John Buckland-Nicks, 1998; Marks, Biermann, Eanes, & Kryvi, 2008; Sasakawa, 2009; Swallow & Wilkinson, 2002; Tavares-Bastos, Teixeira, Colli, & Báo, 2002). In many independently evolved cases, males consistently produce two different sperm types, a phenomenon known as sperm dimorphism. In all cases examined, only one of the two sperm morphs is capable of fertilization (Bressac et al., 1991; Carcupino, Baldacci, Fausto, Scapigliati, & Mazzini, 1999; Eckelbarger, Young, & Cameron, 1989; Sasakawa, 2009; Wilms, 1986). The evolutionary causes and consequences of variation in sperm morphology, both within and between morphs, are immediately intriguing. As gametes, these cells are the final step in the long chain of events leading to reproductive success or failure. Why should such important components of fitness be so variable?

Much of this morphological diversity within morphs can be attributed to deleterious variation, *e.g.* genetic defects (Chenoweth, 2005) or age-related decline in sperm quality (Preston, Saint Jalme, Hingrat, Lacroix, & Sorci, 2015). This deleterious variation has been shown to be inversely correlated with rates of sperm competition between species; taxa that experience more sperm competition tend to have less morphologically variable sperm at both population and individual levels (Kleven, Laskemoen, Fossøy, Robertson, & Lifjeld, 2008). In other words, sperm often vary *in spite of* constraint imposed by their reproductive importance. In species with high rates of polyandry, postcopulatory selection through sperm competition and cryptic female choice weeds out the suboptimal sperm variants, at least for fertilizing sperm (Birkhead, 1998; Immler, Calhim, & Birkhead, 2008).

Production of multiple sperm morphs, conversely, is often posited to be adaptive in some way. The very fact that sperm dimorphism has repeatedly evolved suggests that it has some fitness benefit. Most commonly, non-fertilizing sperm in dimorphic systems are proposed to be specialized agents of male-male competition, acting as final combatants in the struggle for reproductive success (J Buckland-Nicks, Bryson, Hart, & Partridge, 2010; John Buckland-Nicks, 1998; Swallow & Wilkinson, 2002). Indeed, some have suggested that sperm dimorphism allows specialization in the non-fertilizing sperm for a competitor-inhibiting function, sometimes called “kamikaze sperm” (Baker & Bellis, 1989). Although this hypothesis has fallen out of favor, it was proposed and mainly evaluated in the context of mammalian sperm (A. Harcourt, 1991; A. H. Harcourt, 1989; Moore, Martin, & Birkhead, 1999), where non-fertilizing sperm are not usually differentiated from fertilizing sperm in a sophisticated way.

One of the most extreme cases of sperm dimorphism occurs in butterflies and moths (Lepidoptera). In nearly all species of this order, males produce both fertilizing (**eupyrene**) sperm and a second type (**apyrene**) that lacks a nucleus and nuclear DNA (Meves, 1902). The function of apyrene sperm is poorly understood, but because it lacks DNA, it is clearly incapable of fertilizing eggs. Nevertheless, it does not appear to be the result of errors in spermatogenesis; apyrene sperm production is hormonally regulated and occurs in a developmentally predictable way, implying a novel gain of function in these insects (Friedlander, 1997). Organismal studies have demonstrated that males can control the ratio of the two sperm types in their ejaculate and typically transfer to females 10 to 20 times as much apyrene sperm as eupyrene sperm, depending in part on the female’s past mating history (Oberhauser, 1988). These observations have led some to suggest that apyrene sperm play a specialized role in sperm competition (Silberglied, Shepherd, & Dickinson, 1984), yet there remain several other competing hypotheses for apyrene sperm function that have not been resolved through organismal observations and experiments (Swallow & Wilkinson, 2002).

Recently, characterizations of the proteins found in lepidopteran sperm has opened a new avenue to assess their evolution and function (Whittington et al., 2017; Whittington, Zhao, Borziak, Walters, & Dorus, 2015). Proteomic studies have revealed distinct protein profiles for these two cell types (Whittington, Karr, Mongue, Walters, & Dorus, in press). In both morphs, these proteins are retained through maturation, and, in the case of apyrene sperm, the discarding of the nucleus. Because distinct cellular functions are ultimately the product of their expressed protein complement, the class of proteins uniquely found in apyrene sperm make logical targets for understanding the function of these cells from a molecular perspective.

At the molecular level, sperm and other reproductive proteins are often observed to evolve rapidly (Civetta & Singh, 1995; Dorus, Evans, Wyckoff, Sun, & Lahn, 2004; Willie J. Swanson & Vacquier, 2002). For certain reproductive proteins, like sperm-egg interaction pairs, there is compelling evidence that adaptive co-evolution drives this accelerated change (Herberg, Gert, Schleiffer, & Pauli, 2018; W J Swanson & Vacquier, 1998). Yet there are also many instances of reproductive proteins that diverge quickly because of relaxed purifying selection owing to expression in a single sex instead of the whole population (Barker, Demuth, & Wade, 2005; Wade, Priest, & Cruickshank, 2008). Many other factors, including number of protein-protein interactions or importance of reproductive role, can also act to shape the intensity of positive or purifying selection on reproductive proteins (Schumacher, Rosenkranz, & Herlyn, 2014; Schumacher, Zischler, & Herlyn, 2017). Recent theoretical work has formalized the prediction that strong purifying selection on sperm proteins should depend on high rates of polyandry to generate sperm competition (Dapper & Wade, 2016). Thus, with the appropriate datasets, the degree of each sperm morph’s role in sperm competition can be assessed via molecular tests of evolution.

In this study, we report the first molecular evolutionary analyses of dimorphic sperm. We assessed patterns of both polymorphism and divergence among sperm proteins from both eupyrene and apyrene sperm using proteomic datasets of two species: the monarch butterfly, *Danaus plexippus*, and the Carolina sphinx moth, *Manduca sexta* (Whittington et al., in press). North American monarchs spend time at incredibly high density in overwintering colonies in Mexico and California (Urquhart, 1976) and, owing to these unique population dynamics, have some of the highest female remating rates observed in Lepidoptera. Female monarchs mate an average of 2.6 times (and up to 14 times) in overwintering colonies in the wild (Hill Jr., Wenner, & Wells, 1976; Smith, 1984), creating ample opportunity for sperm competition. In contrast, Carolina sphinx moths are typically monandrous (Snow et al., 1974), making sperm competition rarely relevant as a selective force. Taking advantage of this contrast, we investigate the differences in patterns of selection between the two sperm morphs in each species to assess the role of apyrene sperm in sperm competition. If apyrene sperm are involved in sperm competition, their proteins should show evidence of stronger purifying selection in the monarch butterfly. To complete these analyses, we have generated the first published set of whole-genome resequencing data for *Manduca sexta* from a wild population. To test the general predictions for relaxed selection in sex-limited proteins, we used RNA-seq gene expression datasets from previously published data for Carolina sphinx moths (Cao & Jiang, 2017) and newly generated data for the monarch butterfly.

## Materials and Methods

### Sources of data

We used gene sets from the published genomes of each species (Kanost et al., 2016; Zhan & Reppert, 2013) with sperm genes identified from their respective proteomes (Whittington, Karr, Mongue, Walters, & Dorus, in press). We inferred selection from patterns of polymorphism and divergence from congeners using whole genome Illumina resequencing data for both species: a previously published dataset for North American monarch butterflies (Zhan et al., 2014) and a new dataset of North Carolinian sphinx moths. Focal moths were collected with a mercury vapor light trap in July of 2017 in Rocky Mount, North Carolina (see supplemental table S1 for sequencing summary statistics and accessions). Divergences were called by comparison to the queen butterfly (*Danaus gilippus*, previously published in Zhan et al. (2014)) for monarchs, and the five-spotted hawkmoth *(Manduca quinquemaculata*, sequenced for this project) for the Carolina sphinx moth.

In both focal species, we used twelve wild-caught individuals for sampling of polymorphism. In the case of Carolina sphinx moths, these were twelve males caught over the course of three nights. The sex-biased sampling reflects a sex bias in dispersal and collection at the light trap. In the case of monarchs, samples were selected based on depth of sequencing coverage in the published dataset and included 8 females and 4 males from the panmictic North American migratory population. This mixed-sex sampling added the complication of unequal sampling between the autosomes (n = 24) and Z sex chromosome (n = 16). Despite the male-biased gene accumulation on the Z chromosome, the vast majority of sperm genes (92% in the Carolina sphinx, 90% in the monarch) are autosomal in both species (Mongue & Walters, 2017). Due to the sampling complication and limited inference to be gained from Z-linked genes, we focused on the autosomal genes in both species in subsequent analyses.

### SNP-based methods

We aligned sequenced reads with bowtie2 (Langmead & Salzberg, 2012) for conspecifics to their reference genome or with stampy (Lunter & Goodson, 2011) with an increased allowance for substitution for heterospecific alignments. Alignments were taken through GATK’s best practices pipeline (McKenna et al., 2010), including hard filtering, to yield a set of high quality variants both within and between species. Effect-class of each variable site (synonymous, non-synonymous, intergenic, etc.) was determined using custom databases for the two species created with SnpEff (Cingolani et al., 2012). Annotated SNPs were curated to remove false divergences (ancestral polymorphism) and then differences in adaptive evolution were calculated using an estimator of the neutrality index to calculate α, the proportion of substitutions driven by adaptive evolution (Stoletzki & Eyre-Walker, 2011). This form of α corrects the inherent bias in a ratio of ratios while also allowing summation across multiple genes to reduce noise associated with small numbers in count data. For any set of *i* genes with non-zero counts of synonymous (s) polymorphism (P) and divergence (D):

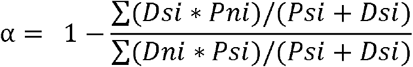

This statistic was calculated with custom scripts in R (R Core Team, 2017).

### Assessment of adaptive evolution and statistical significance

In each analysis, we calculated α for a biologically meaningful set of genes, *e.g.* the sperm proteome and the background genome, and generated a test statistic from the absolute difference of the two point-estimates. To determine significance, we combined the two sets and randomly assigned genes into two new sets of sizes equal to the originals. The difference of these two datasets was determined and the process was repeated for 50,000 permutations to build a distribution of differences between the point estimates of two gene sets of these relative sizes. The p-value was taken as the proportion of times a greater absolute difference was observed between the two random data sets than between the original sets.

We used this permutation approach to make within-species comparisons of α for several different groupings of genes. We first examined differences between the whole sperm proteome and background genome (i.e. all autosomal non-sperm proteins). Next, we considered differences between sperm homologs and sperm proteins unique to one species to assess how selection acted on the same genes in different species. We identified sperm homologs as predicted orthologs that are present in the sperm of both species, with orthology predicted via the proteinOrtho pipeline, as previously reported in Mongue & Walters (2017). Unique sperm proteins may or may not have an ortholog in the other species but are present in the sperm of only one species. Finally, we compared among proteins grouped by their presence in apyrene versus eupyrene sperm. To do so, we classified sperm proteins into three subsets: specific to eupyrene sperm, specific to apyrene sperm, or shared in both types. Pairwise comparisons were made between each subset. For these analyses, we did not consider orthology status owing to the reduction in power that would accompany multiple layers of subdivision of the dataset. For the whole proteome and morph subset comparisons, we further assessed the relative contributions of synonymous and non-synonymous polymorphism and divergence to the α calculation, using a Wilcoxon-Mann-Whitney test to assess significant differences.

### Site-frequency-based methods

We also investigated molecular evolution by leveraging site-frequency-spectrum-based approaches as complimentary evidence. Owing to the redundancy in results, we have included these analyses in the supplement rather the main text. In brief, we used the population genetics software suite ANGSD (Korneliussen, Albrechtsen, & Nielsen, 2014) to generate site frequency spectra at putatively neutral (four-fold degenerate) and selected (zero-fold-degenerate) sites in the genome. We unfolded site frequency spectra and analyzed these spectra with the software polyDFE (Tataru, Mollion, Glémin, & Bataillon, 2017) to examine rates of adaptive evolution in the whole sperm proteomes and background genomes with a more complex likelihood model that corrects for effects of demography and potential misattribution of ancestral state.

### Investigation of sex-limited and tissue-specific expression

Next, we used RNA-seq data to assess whether or not differences in tissue specificity of expression impacted our results from the sperm proteomes in these taxa. For *Manduca sexta*, there already existed a wealth of tissue-specific data at multiple developmental timepoints (Cao & Jiang, 2017). Because we were primarily interested in sperm involvement, we focused on data from adult males, specifically RNA from the testes, head, thorax, and gut. Expression (measured as fragments per kilobase of transcript per million mapped reads, FPKM) was averaged across biological replicates where available in this species. Monarchs had no comparable published data, so we generated separate RNA-seq data sets from the head, thorax, gut, testes, and accessory gland of three adult males (summarized in Table S2 with accessions).

We quantified tissue-specificity of expression using the *specificity metric* (SPM) statistic, a ratio ranging from 0 to 1 indicating the proportion of gene expression occurring in a given focal tissue (Kryuchkova-Mostacci & Robinson-Rechavi, 2017). For instance, a gene with SPM = 0.8 for the testes shows 80% of its total expression across all sampled tissues in the testes. This same gene would have a much lower SPM value in head, thorax, or other tissues. We observed a bimodal distribution of tissue specificities, which allowed us to bin genes into one of two classes: those that displayed low levels of specificity (SPM < 0.5) and those that displayed high levels (SPM > 0.5). After separating genes by specificity, we calculated α for three classes of genes in these two specificity bins.

We had two goals with these analyses: (1) to determine if patterns of adaptive evolution between classes remained the same at both low- and high-specificities and (2) if α increased within a class of genes at higher specificity compared to low. First, we considered background genome genes (i.e. non-sperm genes) ranked by maximum specificity observed in the head, thorax, or gut for each of these genes. Next, we considered only genes identified in the sperm proteome and ranked them by SPM in the testes. Finally, for putatively male-limited non-sperm genes, we excluded sperm proteome genes and considered again those ranked by specificity in the testes (or testes and accessory glands for monarchs). As with our other α calculations, we used non-parametric bootstrapping to generate 95% confidence intervals. For cases in which confidence intervals overlapped, we assessed significance with permutation testing. These analyses were completed with custom R scripts.

### Demographic estimates

Finally, to contextualize the previous analyses and take full advantage of our newly-generated data, we characterized present and historical population sizes of our study species from genomic data. Using folded four-fold degenerate site frequency spectra, we estimated neutral coalescence patterns with Stairway Plot (Liu & Fu, 2015). For estimated generation time, we used four generations per year for monarchs and three for the Carolina sphinx moth. For mutation rate, we chose the estimate 2.9*10^−9^ from the butterfly *Heliconius melpomene*, the closest relative with a spontaneous mutation rate estimate (Keightley et al., 2015).

## Results

### Differences Between Sperm Proteins and the Background Genome

First, we considered the sperm proteome as a whole (*i.e.* all apyrene, shared, and eupyrene proteins) and compared adaptive evolution of genes found in sperm to those in the background genome, defined as all autosomal protein coding genes not present in the sperm proteome. Z-linked genes were excluded from the analysis. We counted and classified synonymous and non-synonymous single nucleotide polymorphisms within species and divergences to a congener (*Danaus gilippus* for the monarch, and *Manduca quinquemaculata* for the Carolina sphinx). These quantities were used to generate an estimate of the proportion of adaptive substitutions (α) per gene-class for both the sperm proteome and the background genome. We found no difference in α between the sperm proteome and the rest of the genome in the Carolina sphinx (p = 0.40892 by permutation testing, Figure 1A, left); for monarchs, however, the sperm proteome showed a significantly greater proportion of adaptive substitutions than the rest of the genome (p = 0.00006, Figure 1A, right). Note that in the strict sense, negative α values are not biologically meaningful and likely point to an abundance of weakly deleterious variants within populations or complex demographic histories (Eyre-Walker & Keightley, 2009); nevertheless, these confounding variables should not differentially affect genes within species, so our observed differences point to true differences in selection in gene sets.

**Figure 1.**
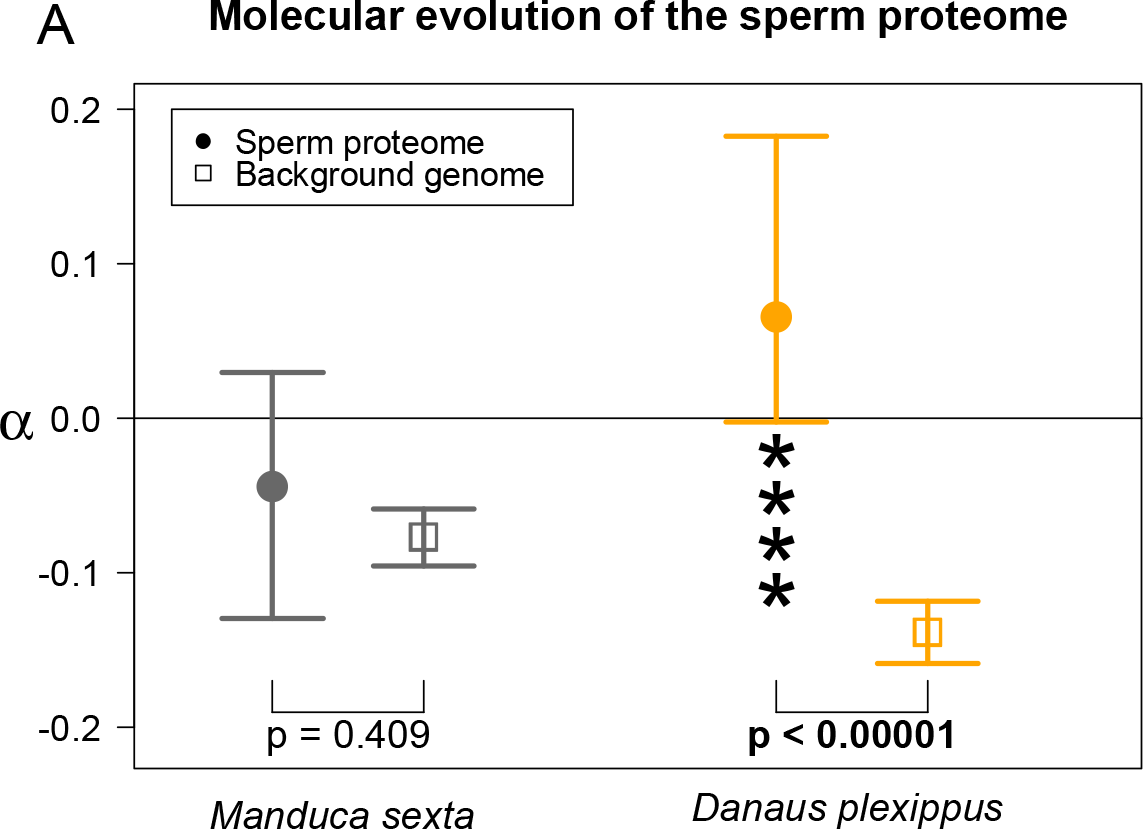
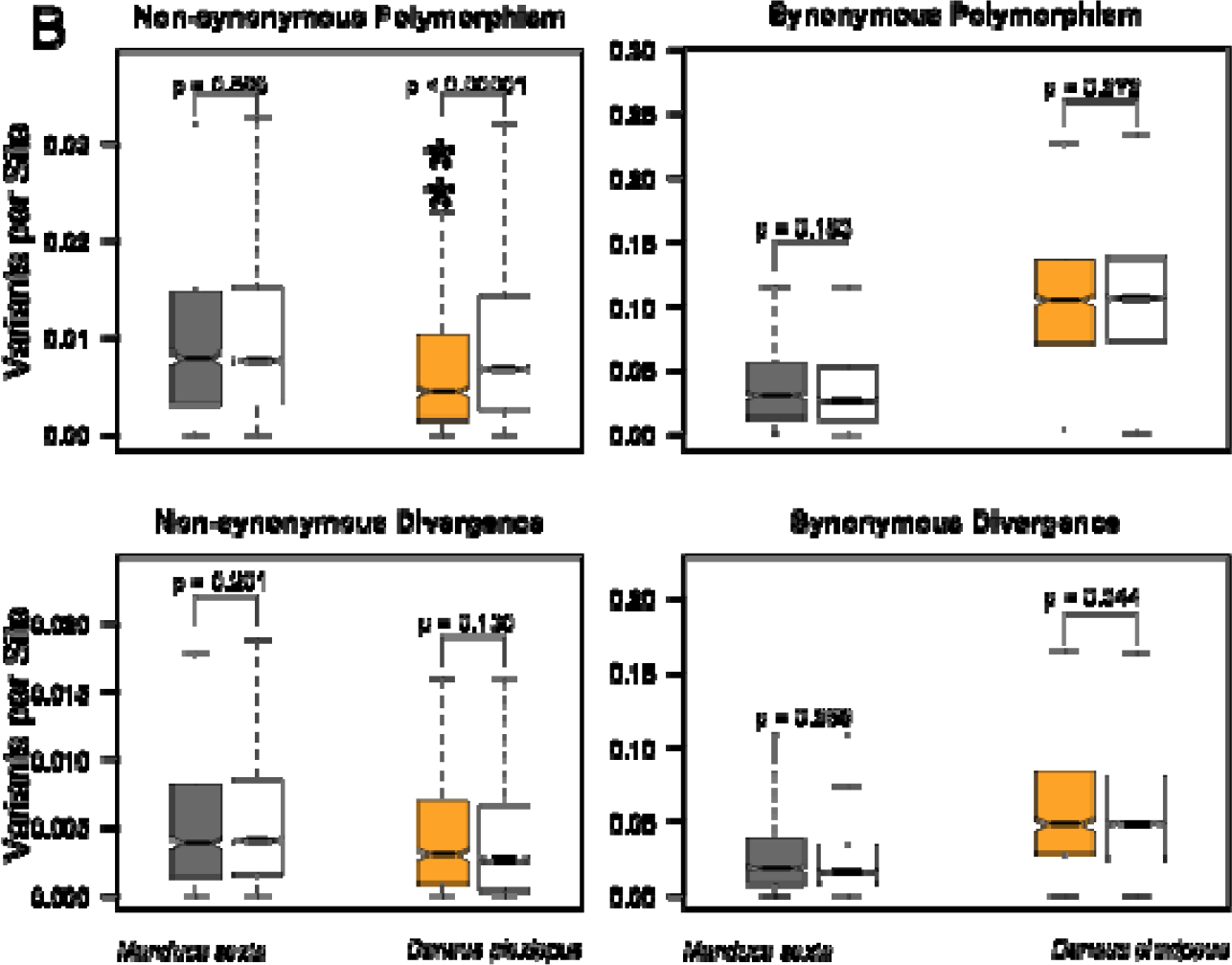
**A.** In the Carolina sphinx moth (*M. sexta*), there is no difference between the sperm proteome and the rest of the genome (left); conversely, genes in the sperm proteome of monarch butterflies (*D. plexippus*) show a significantly higher proportion of adaptive substitutions (α) than the rest of the genome (right). P-values come from permutation tests. Error bars represent 95% bootstrapped confidence intervals from the point estimates. **B.** Decomposing α into its components: Pn, Ps, Pn, and Ds and comparing the sperm proteome (filled boxes) to the background genome (open boxes). There were no strong differences between sperm genes and the background genome in Carolina sphinx moths. In monarch butterflies, the signal for increased adaptive substitution comes from a marginal increase in non-synonymous divergence (bottom left) combined with a great reduction in non-synonymous polymorphism in sperm genes compared to the rest of the genome (top left). P-values reflect Wilcoxon-Mann-Whitney tests, with * < 0.05, ** < 0.005, *** < 0.0005, etc.

To better understand the relative roles of polymorphism and divergence in sperm and background genes, we investigated the individual components of α: counts of non-synonymous polymorphism (Pn), synonymous polymorphism (Ps), non-synonymous divergence (Dn), and synonymous divergence (Ds). We compared the scaled estimates of each (*e.g.* non-synonymous polymorphisms per non-synonymous site) to the background genome within each species using a Wilcoxon-Mann-Whitney test (Figure 1B).

We found no differences between sperm and the background for any class of variants in *M. sexta* (Pn: W = 3014100, p = 0.5964; Ps: W = 2879300, p = 0.1830; Dn: W = 3068300, p = 0.2009; Ds: W = 2895700, p = 0.2686). The signal for elevated α in monarch sperm primarily reflects non-synonymous polymorphism, which was greatly depressed (W = 3062400; p = 3.224 * 10^−11^), as would be expected under strong purifying selection, while other classes were comparable between sperm and the background genome (Ps: W = 2684200, p = 0.2720; Dn: W = 2506400, p = 0.1300; Ds: W = 2544400, p = 0.3437).

Next, we leveraged orthology, as established by Whittington *et al.* (2017), to test for differences in mating system while controlling for the effects of sperm proteome content. Substantial numbers of orthologous proteins are found in the sperm proteomes of both species, which we hereafter referred to as *sperm homologs*. Sperm homologs offer the opportunity to directly assess the selective pressures experienced by the same genes with putatively conserved function but found in species with different levels of postcopulatory selection. Nearly half of the monarch sperm proteome (^~^42%, 216 genes, Figure 2A) shares an ortholog in the sperm proteome of *M. sexta*; reciprocally, there are 236 genes (37%) in the Carolina sphinx sperm proteome that share an ortholog in the monarch sperm proteome; these numbers are not equal due to lineage-specific duplications among sperm homologs creating a few cases of one-to-many orthology. We tested for differences in adaptive evolution between sperm homologs and sperm proteins unique to one species (orthology outside of sperm or no detectable orthology). In Carolina sphinx moths, genes of these two classes did not differ in the proportion of adaptive substitutions with permutation testing (p = 0.6174, Figure 2B). In monarchs, we detected an increased proportion of adaptive substitution in the sperm homologs compared to unique proteins (p = 0.0372, Figure 2B). Comparing between species, sperm homologs had much higher α values in monarchs than in Carolina sphinx moths (p = 0.00008), while genes with unique expression in either species did not show differences between species (p = 0.5922). Thus, the same sperm proteins appear to be evolving more adaptively in the polyandrous species.

**Figure 2.**
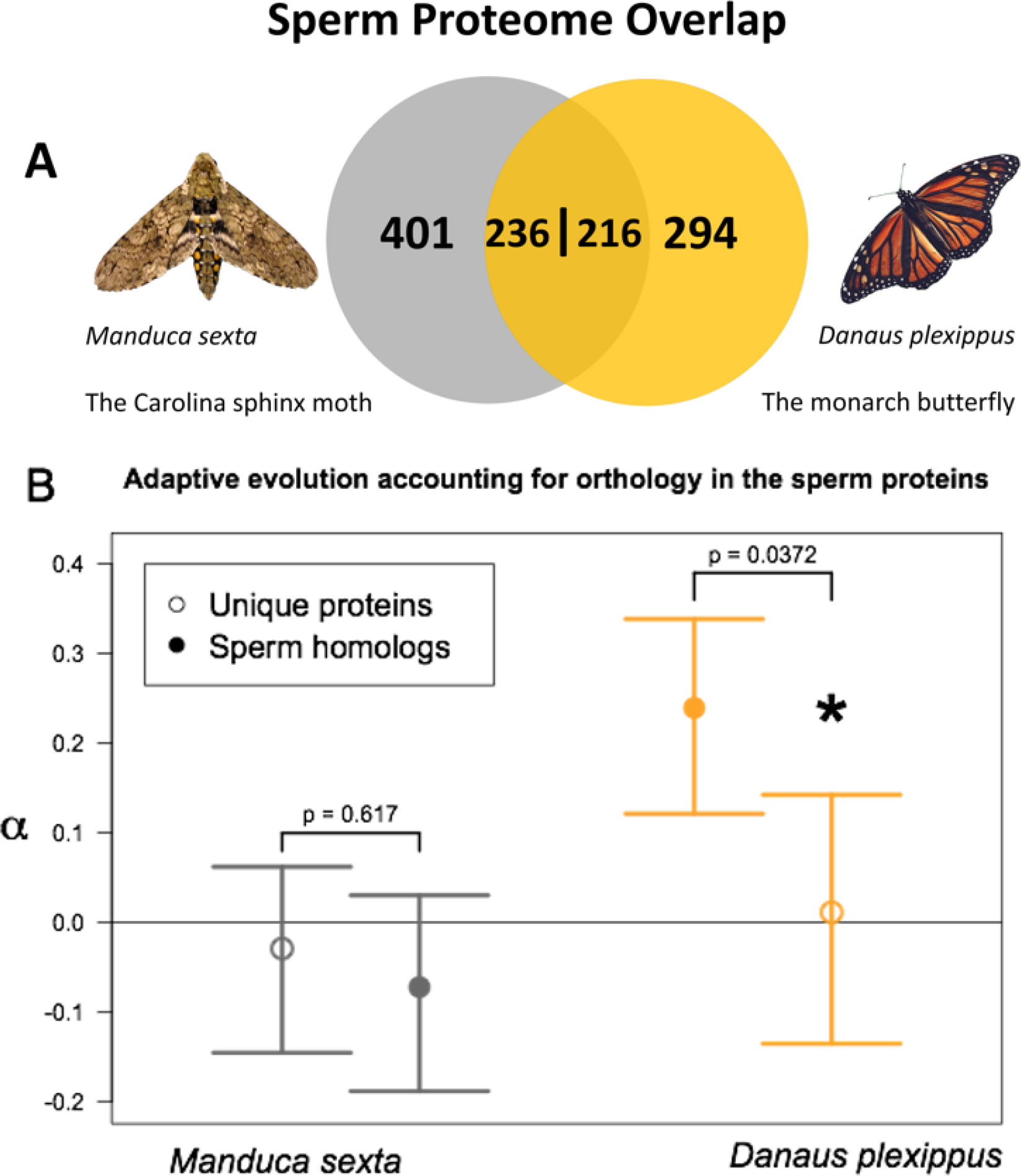
**A.** Composition of the portion of the sperm proteomes analyzed in this study. Numbers indicate counts of proteins unique to one species’ sperm or with an ortholog in the other species’ sperm (sperm homologs). Note that the overlap number varies between species due to the presence of a few one-to-many-orthologs. **B.** Sperm homologs show evidence for a greater proportion of adaptive substitutions (α) in monarch butterflies, but not in Carolina sphinx moths. P-values are based on permutation tests comparing the difference between two sets of genes randomly assigned from the sperm proteome in each species; error bars are 95% bootstrap confidence intervals.

### Site-frequency based methods

We also took a likelihood approach to modeling adaptive evolution using site frequency spectra generated from the same samples we used forSNP-counting. These results are detailed in the supplement. In short though, we found a shift in the predicted distribution of fitness effects of new mutations in monarch sperm proteins compared to the background consistent with stronger purifying selection (Figure S1) and drastically higher α in sperm genes in monarchs alone (Figure S2).

### Patterns of Adaptive Evolution in Sex-Specific Tissues

Next, we used RNA-seq data to examine the effect of tissue-specificity on selection in these insects. With these data, we calculated the tissue specificity metric, SPM (Kryuchkova-Mostacci & Robinson-Rechavi, 2017), which ranges from ubiquitous expression (near 0) to single-tissue specific (1). Although the sperm proteomes of both of our species were enriched for gene products specifically expressed in testes, they also contained broadly expressed gene products (Figure 3B). To assess the effect of these broadly expressed genes on our inference of selection, we recalculated the α statistic for two bins of genes (Figure 3C): those with broad expression (SPM < 0.5) and those with high tissue-specificity (SPM > 0.5).

**Figure 3.**
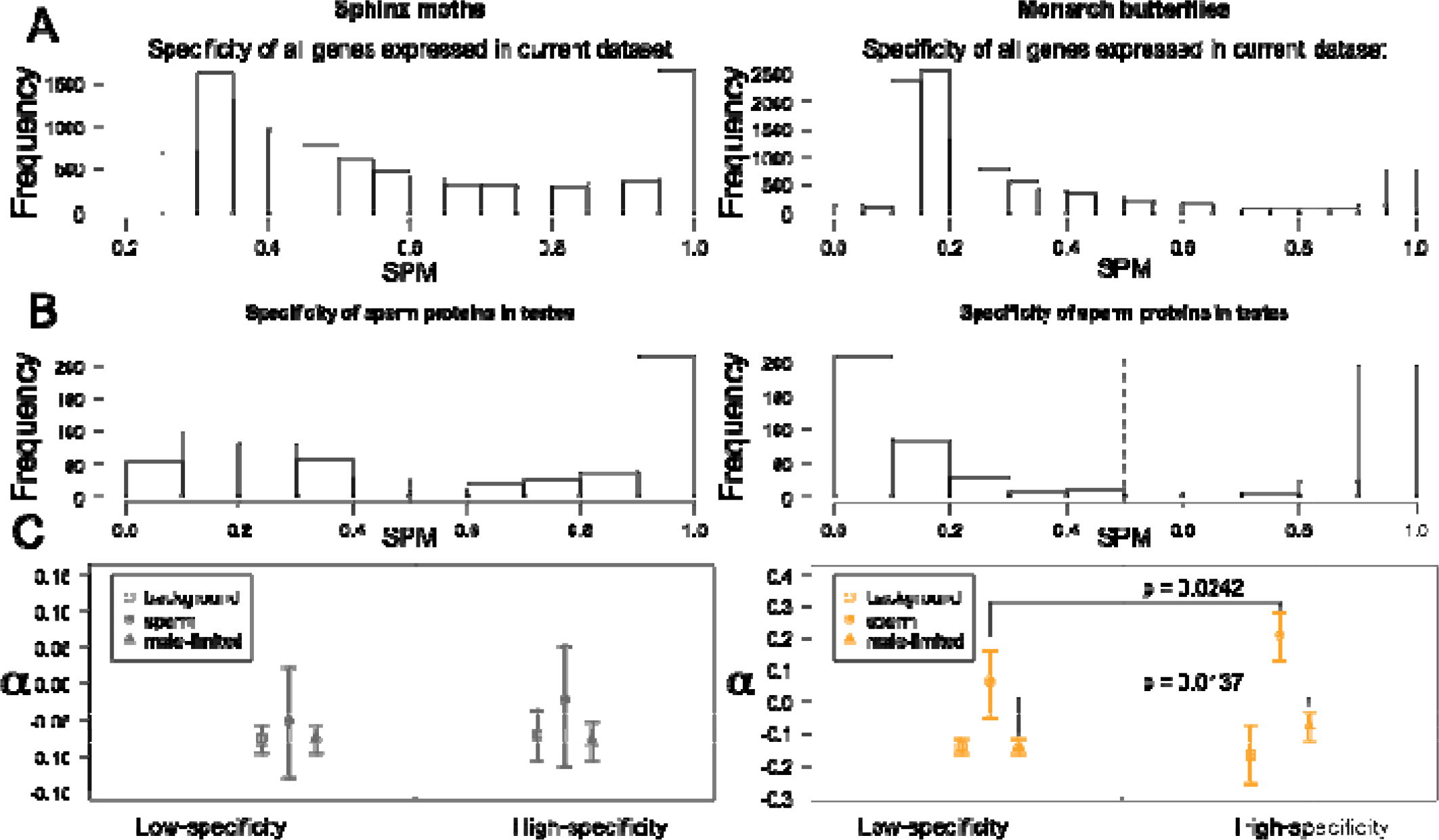
Investigating how tissue specificity of gene expression impacts adaptive evolution in both Carolina sphinx moths (**left column**) and monarchs (**right column)**. **A.** Maximum specificity of all genes across all studied tissues using the RNA-seq data considered in these analyses. **B.** Observed distribution of specificity of sperm proteome genes expressed in the testes. Based on these distributions, we separated genes into one of two categories, low-specificity (SPM < 0.5) or high-specificity (SPM > 0.5), divided by the dashed line. **C.** Inferred proportion of adaptive substitutions (α) in background genes (squares), sperm proteome genes (circles), and male-limited genes (as defined by testes expression). Bars represent 95% confidence intervals from non-parametric bootstrapping. Non-overlapping confidence intervals imply significant differences generally, but we have also highlighted two significant differences that are less visibly apparent. Monarchs show evidence for increasing α with increasing tissue specificity in sperm and testes genes, but sphinx moths do not. Moreover, sperm proteome genes evolve more adaptively than background or testes-specific genes in both specificity groups for monarchs but not sphinx moths.

In Carolina sphinx moths, there were no significant changes in α between low- and high-specificity genes in any part of the genome (background genes: p = 0.3868, sperm proteome genes: p = 0.3248, male-limited genes: p = 0.5579; Figure 3C, left), nor did any of the gene classes differ from each other within a specificity bin. In monarchs, however, both sperm proteome genes (p = 0.0242) and testes genes (p = 0.0137) showed higher α in the high-specificity group than the low-specificity group, though somatically expressed genes did not (p = 0.6831). Additionally, we found that sperm genes showed much greater α than the background genome or other genes expressed in the testes at both low- and high-specificities (Figure 3C, right). This result indicates that our initial results (considering the whole sperm proteome) are not dependent on the underlying specificity of sperm genes.

### Molecular evolution in dimorphic sperm

Having verified the patterns of evolution in the whole sperm proteomes with several approaches, we turned to our primary question, assessing apyrene sperm function through analysis of molecular evolution. We considered the different subsets of the sperm proteomes based on the two sperm types. The two datasets consisted of three classes of sperm proteins: unique to eupyrene sperm, unique to apyrene sperm, or found in both cell types (henceforth “shared”, Figure 4A). We assessed differences in selective pressures between the sperm morphs with another series of permutation tests, both comparing parts of the sperm proteome to the background genome and comparing parts of the proteome to each other.

**Figure 4.**
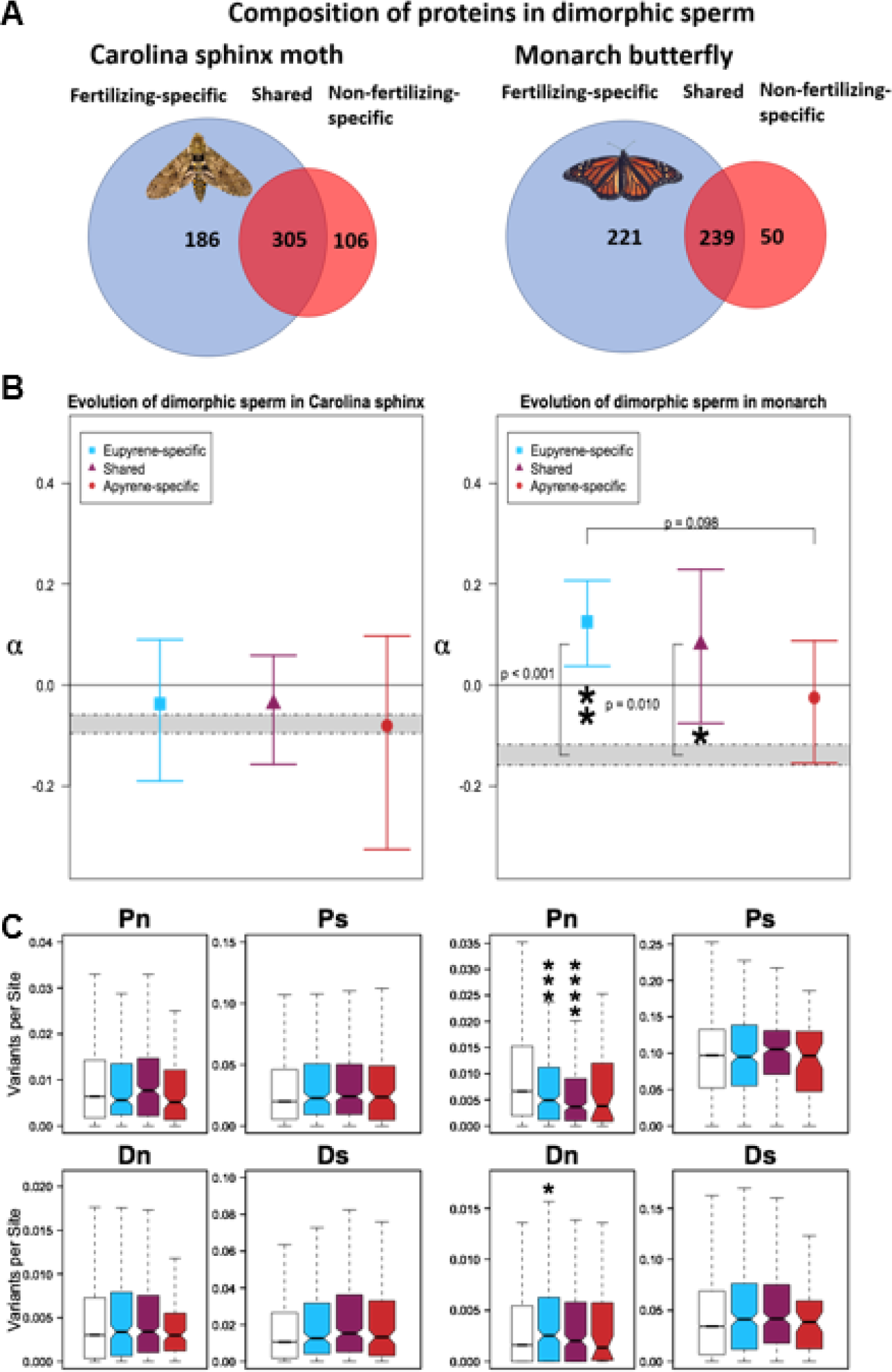
**A.** Composition of the sperm proteome with respect to dimorphic sperm. The majority of identified proteins were shared between the two cell types, followed by the set unique to eupyrene sperm, and finally the smallest set was the proteins found only in apyrene sperm. **B.** None of the sets of sperm proteins evolved either differently from each other or distinctly from the background genome (shaded regions represent 95% confidence intervals of the background) in the Carolina sphinx (left). In the monarch however (right), the signal for elevated α was localized to the eupyrene-specific and shared proteins. There was also a trend for increased α in eupyrene-specific proteins as compared to apyrene-specific. Error bars represent 95% confidence intervals from bootstrapping. **C.** Decomposing α into Pn, Ps, Dn, and Ds for dimorphic sperm. Plotting of variation follows the coloring and order in parts A and B; from left to right in each panel: background genome, eupyrene, shared, and apyrene sperm. Asterisks denote significant differences from the background genome based on a Wilcoxon-Mann-Whitney test, with * < 0.05, ** < 0.005, *** < 0.0005, etc.

As expected based on the whole-proteome results from Carolina sphinx moth, neither eupyrene-specific (p = 0.55912), shared (p = 0.4647), nor apyrene-specific proteins (p = 0.96496) differed from the background genome (Figure 4B). α did not vary between apyrene-specific and eupyrene-specific proteins (p = 0.7271), between apyrene-specific and shared (p = 0.7176) or eupyrene-specific and shared proteins (p = 0.9979). In monarchs, both eupyrene-specific proteins (p =0.00018) and shared proteins (p = 0.01038) showed elevated α, but apyrene-specific proteins did not evolve differently from the background genome (p = 0.55934). Neither apyrene nor eupyrene sperm differed significantly from the shared set in monarchs (p = 0.6332 & p = 0.6234, respectively), but there was a trend towards significantly increased α in eupyrene-specific proteins compared to apyrene-specific proteins (p = 0.0986).

As with the whole sperm proteome, we investigated which classes of variants contributed to our observed differences in α (Figure 4C). Consistent with the results above, none of the variant classes significantly differed from the genome background in the sphinx moth eupyrene-specific proteins (Pn: W = 995190, p = 0.0857; Ps: W = 943550, p = 0.6782; Dn: W = 966630, p = 0.3183; Ds: W = 963410, p = 0.3596). Shared proteins also showed the same level of variation as the background across all variants (Pn: W = 1470300, p = 0.3277; Ps: W = 1444400, p = 0.1369; Dn: W = 1540700, p = 0.6883; Ds: W = 1437100, p = 0.1030). And finally, apyrene-specific proteins were not significantly different either (Pn: W = 548570, p = 0.4974; Ps: W = 491410, p = 0.2149; Dn: W = 560910, p = 0.2741; Ds: W = 495180, p = 0.2653). In summary, there was no evidence for stronger selection on either sperm morph in Carolina sphinx moths.

For monarchs, we found that the elevated α in the eupyrene-specific and shared subsets was driven primarily by a decrease in non-synonymous polymorphism compared to the background genome (W = 1291700, p = 0.0003 for eupyrene; W = 1486100, p = 1.167*10^−8^ for shared). Apyrene-specific proteins did not show a reduction in non-synonymous polymorphism (W = 284620, p = 0.1684). Synonymous polymorphism did not significantly differ from the background in any subset of the sperm proteome (eupyrene: W = 1164200, p = 0.4492; shared: W = 1249900, p = 0.5570, apyrene: W = 270160, p = 0.4927). Nor did synonymous divergence (eupyrene: W = 1056000, p = 0.0928; shared: W = 1209000, p = 0.7665, apyrene: W = 279420, p = 0.2594). Intriguingly, non-synonymous divergence was elevated compared to the background in eupyrene-specific proteins (W = 1021800, p = 0.0151), but not the shared (W = 121800, p = 0.9185) or apyrene-specific portions of the proteome (W = 266580, p = 0.6042). This suggests periodic sweeps of positively selected variants in fertilizing sperm proteins.

We did not examine orthology within dimorphic sperm owing to small gene counts giving reduced statistical power. Nor could we could examine tissue specificity here because apyrene and eupyrene sperm are produced at different developmental timepoints and we did not have suitable expression data in both species. Nonetheless, the consistency of results in the whole proteome datasets gives us no reason to expect that within-proteome results would be idiosyncratic to our methodology.

### Demographic estimates

Finally, to contextualize our results with population dynamics, we estimated population size history using site frequency from 4-fold degenerate sites in the two species’ genomes (Figure S3). Both have effective population sizes near 2,000,000, as expected of herbivorous invertebrates with high dispersal potential, numerous host plants, and a large range over North America. We also recovered a population size increase in monarch butterflies in the recent past, which has been previously reported with genomic data (Zhan et al., 2014). We note that our inferred timing of this event differs from that of the previous authors, who used mutation rate estimates from *Drosophila melanogaster.* Such input parameter differences affect the estimated time of events, but not the trajectories.

## Discussion

We investigated the molecular evolution of eupyrene (fertilizing) and apyrene (non-fertilizing) sperm, the ubiquitous lepidopteran cell-type of unknown functional significance. These sperm have long been posited to interfere with competitors’ sperm, in part because their quantity varies with levels of male-male competition (Silberglied et al., 1984; Solensky & Oberhauser, 2009; Swallow & Wilkinson, 2002). In contrast to these organismal observations, the results of our molecular analyses cast doubt on this hypothesis. If apyrene sperm played an active role in sperm competition, we would expect evidence for stronger selection in apyrene sperm compared to the background genome in monarchs. We found a signal for elevated adaptive evolution (α) in the sperm proteome compared to the background genome in these polyandrous butterflies, but this signal did *not* include apyrene-sperm-specific proteins. Instead, genes encoding apyrene sperm proteins evolve similarly to the background genome in both monarchs and Carolina sphinx moths. This result is unlikely to have arisen from a lack of power in our methodologies, as eupyrene-specific and shared sperm proteins showed patterns in line with expectations for a role of sperm competition in molecular evolution in monarchs.

### Selection consistent with sperm competition, but only in fertilizing sperm

The source of the apparently elevated α in the monarch sperm proteome came mainly from a dearth of non-synonymous polymorphisms in sperm proteins compared to the background genome, indicating the action of purifying selection to remove many variants before fixation in monarchs. Strong purifying selection has been similarly observed in genes expressed in pollen, the main male-male competitors in flowering plants (Arunkumar, Josephs, Williamson, & Wright, 2013). A similar pattern can also be observed in passerine birds, in which species with higher rates of sperm competition show less intraspecific and intra-male variation in sperm length compared to sperm of less polyandrous species (Immler et al., 2008; Kleven et al., 2008).

Moreover, the elevated α in sperm homologs in monarchs suggests that genes that have had conserved sperm function since the divergence of the two species some 100 million years ago (Heikkila, Kaila, Mutanen, Pena, & Wahlberg, 2012) are under stronger purifying selection in the polyandrous species. According to recent gene ontology analyses, such genes are enriched for core traits in sperm, such as mitochondrial function, respiration, and flagellar structure. Similarly, proteins shared between the two sperm types and those unique to eupyrene sperm show an elevated α compared to the background genome in monarchs. Sperm proteins shared between morphs are enriched for structural proteins that give rise to the sperm tail and thus impact motility (Whittington et al., in press), while those expressed only in eupyrene sperm doubtless include important mediators of fertilization. At the cellular level, variation in sperm traits like swimming ability, longevity, and overall viability affects sperm competition outcomes (Burness, Casselman, Schulte-Hostedde, Moyes, & Montgomerie, 2004; Kim et al., 2017) and has a polygenic basis in other taxa (Hering, Olenski, & Kaminski, 2014). For traits like longevity and motility there is a threshold below which fertilization becomes significantly impaired, but in the absence of competitor alleles, there is a larger range of effectively-neutral trait-values, allowing for more variation to be maintained in the population. In the presence of competitor alleles, however, marginal differences in fertilization success come under selection, leading to the removal of deleterious variants through sperm competition.

Stronger selection from competition may include even the event of fertilization itself. Lepidopteran eggs are known to possess multiple micropyle openings for sperm (Kumar, Kariappa, Babu, & Dandin, 2007) and eupyrene sperm possess structures resembling an acrosome (while their apyrene counterparts do not) (Friedlander, 1997). This rare combination of male and female gamete structures is also found in sturgeon, in which the multiple micropyles give several sperm potential access to the egg nucleus and there is competition among sperm to initiate karyogamy via the acrosome reaction (Psenicka, Rodina, & Linhart, 2010). Consistent with micropyle-mediated competition, it has been shown that more polyandrous species of Lepidoptera tend to have more micropyles on their egg surfaces than monandrous species (Iossa, Gage, & Eady, 2016). If this truly does extend the opportunity for male-male competition and cryptic choice, then acrosomal proteins in eupyrene sperm would be likely targets for selection in polyandrous systems.

Whatever the mechanics of fertilization are, paternity outcomes in polyandrous species are often bimodally distributed (Simmons & Siva-Jothy, 1998; Wedell & Cook, 1998), including in monarch butterflies (Mongue, Ahmed, Tsai, & De Roode, 2015). For females that mate twice, one of the two males typically fathers most, if not all, of the observed offspring produced by the female, but there is little consistency in whether it is the first or second male. With these dynamics, fitness differences between winning and losing sperm phenotypes are large and selection can reliably remove less successful genotypes.

Evidence of this can be seen in the estimated distribution of fitness effects of new mutations in monarch sperm proteins. Compared to the background genome, we see a decrease in the proportion of effectively neutral and weakly deleterious mutations and an increase in both strongly deleterious and beneficial mutations. In the absence of competition, not only are mildly suboptimal variants effectively neutral, but novel, more efficient competitors should have no selective advantage in monandrous species unless they also markedly increase fitness in a single mating. This reasoning is supported by the estimated distribution of fitness effect for the complimentary gene sets in the Carolina sphinx moth; in this species, we see little variation in the DFE between the background genome and the sperm proteome. Moreover, there is no decrease (and indeed) an increase in non-synonymous divergence of eupyrene sperm proteins in monarchs compared to the rest of the genome. This pattern suggests that in addition to strong purifying selection there must be periodic sweeps of beneficial alleles. Without a broader, phylogenetically controlled study, these results between a single pair of species are not conclusive, but they fit well with the prediction that sperm protein evolution depends on the rates of polyandry in a species (Dapper & Wade, 2016).

### Evolution of tissue-specific and male-limited genes

Other studies have demonstrated that tissue specificity of expression can strongly influence the molecular evolution of reproductive proteins (Schumacher & Herlyn, 2018), in some cases more than mating system (Carnahan-Craig & Jensen-Seaman, 2014). Because our proteomic data did not contain information on tissue specificity on their own, we examined this dynamic with RNA-seq data. We found increased adaptive evolution in monarch sperm genes with higher specificity compared to low-specificity sperm genes. Furthermore, while not significantly different from background genome, α for non-sperm genes expressed in the testes increased with greater specificity in monarchs, suggesting that they too may be subject to stronger sexual selection in this polyandrous species. Neither of these patterns held for Carolina sphinx moths, which showed no differences based on tissue specificity. This consistency further suggests the difference in mating system as an explanation for differences in intensity of selection.

Finally, we did not observe relaxed constraint in reproductive proteins predicted due to the smaller effective population size of males or females compared to the population as a whole, as predicted by theory (Dapper & Wade, 2016; Wade et al., 2008). Specifically, we did not observe a difference in the adaptive evolution of genes with testes-specific expression, our proxy for sex-limited expression, compared to the background genome. To explain this discrepancy between theory and observation, we turn to Nearly Neutral Theory. Large populations have more efficient selection than small populations and a smaller range of slightly deleterious mutations that behave neutrally (Ohta, 1992). Mutations with a selective effect less than 1/N_e_ are expected to behave neutrally. For instance, one commonly cited estimate for human population size is Ne ≈ 10,000 over evolutionary history (Zhao et al., 2000). Based on this, mutations with selective effects less than 0.0001 should behave neutrally for alleles expressed in both sexes, while those with effects of 0.0002 are effectively neutral for alleles only expressed in one sex. And indeed, there is evidence that genes expressed only in men have a higher mutational load than those expressed in both sexes (Gershoni & Pietrokovski, 2014). Chimpanzees, another species with a similar effective population size (Won & Hey, 2005), also show increased non-synonymous divergence in reproductive proteins (Wong, 2010). Broadly, male reproductive protein evolution appears to depend more on effective population sizes than intensity of sperm competition in the great apes in general (Good et al., 2013), as one would expect for species with relatively small effective population sizes.

In contrast to mammals, the effective population sizes of most insect species are orders of magnitude higher. Using neutral site frequency spectra, we estimated effective populations near 2,000,000 for both North American monarchs and Carolina sphinx moths. Selection is much more effective in these massive populations; mutations with effects above 5*10^−7^ should be subject to selection in both sexes and those above 1*10^−6^ should be subject to selection if expression is sex-limited. Thus, even selection on alleles with sex-limited expression in these insects should be 100 times stronger than selection on the entire human population. Even if there is a relative two-fold difference in selection, the absolute magnitude of the difference should be miniscule, and the effects of mating system more apparent.

### Advancing understanding of apyrene sperm

Previous morphological work found that eupyrene sperm traits (like sperm length) but not apyrene sperm traits, varied with risk of sperm competition in other butterflies (Gage, 1994). Similarly, from a molecular perspective, none of the patterns of increased purifying and positive selection that we observed for monarch sperm proteins applied to the apyrene-specific proteins. That we also do not see evidence for the action of sperm competition on apyrene-specific protein evolution is itself informative, however. Research to-date has proposed four main hypotheses for apyrene sperm (Swallow & Wilkinson, 2002): active sperm competition agents, passive competition agents, nutrient nuptial gifts, or necessary facilitators of fertilization. Our molecular analyses argue against apyrene sperm as active agents of sperm competition, but it is worth considering predictions for molecular evolution of apyrene sperm under the other hypotheses.

Indeed, apyrene sperm may still have adaptive significance without specialized molecular function, especially under the filler hypothesis. This proposed function also relates to sperm competition, but posits that apyrene sperm are employed proactively, to fill the female’s sperm storage organ and delay remating, thus decreasing the risk of sperm competition, rather than impacting its outcome (Swallow & Wilkinson, 2002). Both in monarchs and the butterfly *Pieris napi*, female time to remating increases with the number of apyrene sperm received from males (Cook & Wedell, 1999; Oberhauser, 1988). Such observations are somewhat confounded by the size of the spermatophore nuptial gift that males provide during mating, but apyrene sperm themselves have been proposed as a form of nutritional nuptial gift (He, Tanaka, & Miyata, 1995; Lamunyon, 2000). Under both the nutrient and filler hypotheses, the actual sequence of apyrene sperm proteins should be less important than their physical presence and abundance, so factors affecting the rate of apyrene sperm production would be more likely targets for selection in polyandrous species than the proteins sequences themselves.

Finally, apyrene sperm appear to capacitate fertilization in *Bombyx mori* (Takemura, Sahara, Mochida, & Ohnuma, 2006); the mechanism here is unclear and the phenomenon is untested in other taxa, but it could conceivably involve proteins that modulate female reproductive physiology to make conditions more favorable for eupyrene sperm or induce oviposition. In such a case, these proteins would behave more akin to the broader class of reproductive proteins and evolve independently of rates of polyandry in a species. If there is an evolutionarily conserved capacitation effector in our study taxa, it is possible that this function is governed by a small subset of apyrene-specific proteins. Because our methods aggregate signal for selection across multiple genes or sites to counteract high variance in variant counts within genes (Stoletzki & Eyre-Walker, 2011), the importance of one or a few genes could be lost in the heterogeneous selection on different proteins.

## Conclusions

Variation in reproductive traits has long been studied at the morphological and molecular level, generally. Yet sperm dimorphism, one of the most striking and enigmatic reproductive traits, has not previously been assessed using population genetic analyses. Our investigation of the sperm proteome in two Lepidoptera demonstrates a pattern of stronger purifying selection on fertilizing-sperm genes in a species with higher rates of sperm competition. In this polyandrous species, these genes experience a strikingly different selective environment than the rest of the genome, with strong purifying selection reducing variation in sperm genes. In contrast, fertilizing-sperm genes in the monandrous species hold as much deleterious variation as other parts of their genome. Our new molecular findings fit well with established studies on sperm morphology which show that sperm competition results in decreased variation in sperm traits.

The evolution of non-fertilizing sperm, however, does not show a strong influence of sperm competition. This lack of pattern itself argues against apyrene sperm as active agents of sperm competition, one of the long-held hypotheses for non-fertilizing sperm function. Instead, apyrene sperm may play a passive role in reducing the risk of competition by delaying female remating. The method by which apyrene sperm capacitate fertilization in some species remains unclear based solely on genomic approaches and will likely require functional experiments to completely understand.

## Acknowledgments

This project was funded by the NSF DDIG (DEB-1701931) and Kansas Idea Network of Biomedical research (NIH P20 GM103418). The authors wish to acknowledge Wesley Mason and Michael Hulet and the rest of the Information and Telecommunication Technology Center (ITTC) staff at the University of Kansas for their support with our high-performance computing. Thank you to Jacobus de Roode for use of the monarch image, Elizabeth Moore for facilitating collaboration between Kansas and North Carolina, Tawny Scanlan for comments on sperm biology, and Amanda Pierce and Tom de Man for housing during field collection.

## Author Contributions

AJM designed the experiments, collected samples, performed analyses, and wrote the manuscript. MEH collected samples and conducted analyses. LG provided data and performed analyses. CES planned and facilitated sample collection and edited the manuscript. JRW assisted in experiment design and manuscript editing.

## Data accessibility

*Manduca sexta* whole genome resequencing data can be found on NCBI’s Sequence Read Archive with the following accession: SRP144217. *Danaus plexippus* RNA sequencing data can be retrieved with accessions: SRR8580831 – SRR8580842. Analysis scripts can be found at https://github.com/WaltersLab/DimorphicSpermMolEvo.

