## Supplemental Figures for "Molecular evolution of non-fertilizing sperm in Lepidoptera suggests minimal direct involvement in sperm competition"

**Supplemental Materials**

*Site-frequency-based analyses*

Starting with the same monarch and sphinx moth samples used for SNP-calling, we used the population genetics software suite ANGSD (Korneliussen, Albrechtsen, & Nielsen, 2014) to generate site frequency spectra at putatively neutral (four-fold degenerate) and selected (zero-fold-degenerate) sites in the genome. We unfolded site frequency spectra using parsimonious inference of ancestral state of alleles. These unfolded spectra were analyzed with polyDFE (Tataru, Mollion, Glémin, & Bataillon, 2017). This method uses information from four-fold sites to develop an expectation for the distribution of neutral allele frequencies owing to mutation, demographic processes, and error. This neutral distribution is compared to the distribution from sites under selection (zero-fold sites), and the differences between the two distributions can attributed to selection.

We compared sites from the backgrounds to sites from the sperm proteomes to see if estimates of α or the distribution of fitness effects of new mutations (DFEs) differed between these two gene sets in each species. Divergence counts were omitted here to simplify the likelihood computation for these large datasets and remove any error from misattributed divergence. To place confidence intervals on estimates of α and the DFEs, input site frequency spectra were parametrically bootstrapped to obtain 100 simulated site frequency spectra of similar distribution. Processing of model inputs and outputs was accomplished with R scripts.

The DFEs of new non-synonymous mutations suggest stronger selection on sperm genes than the rest of the genome in monarchs but not Carolina sphinx moths. In Carolina sphinx moths, the DFE is quite consistent across the background genome, the whole sperm proteome, and the sperm homologs in the (Figure S1, left panels). By contrast, the DFE in monarchs differs substantially between the background genome and the sperm proteome. Relative to the background, the sperm proteome shows a dearth of weakly deleterious and effectively neutral variants, with a concurrent increase in strongly deleterious and beneficial variants. This pattern is even more exaggerated in the sperm homologs, where almost no neutral variants are detectable (Figure S1, right). Indeed, the decrease in weakly deleterious variants in monarch sperm genes should weaken potential downward bias in the SNP-based α calculation compared to the background (Eyre-Walker & Keightley, 2009b), so the observed difference in α from our previous analysis may be exaggerated. However, the increase in positively selected variants in both the whole sperm proteome and sperm homologs suggests that there is a true increase in adaptive evolution compared to the background genome.

Using these likelihood models provided by polyDFE to estimate α, we see a slight increase for the sphinx moth sperm proteome compared to the background, but this pattern disappears when considering only sperm homologs. Moreover, we see a much larger difference in selection on sperm protein variants compared to the rest of the genome in monarch butterflies. Upwards of 90% of substitutions in monarch sperm proteins are inferred to be a result of adaptive evolution in both the whole sperm proteome and the shared sperm orthologs (Supplemental Figure 1). We note that estimates of α are influenced by the ways in which demography is (or is not) accounted for (Messer & Petrov, 2013), so it is not surprising that the values obtained with this more complex likelihood method differ from our estimates based directly on simple counts of polymorphism and divergence. Likely, the values estimated via likelihood modeling are closer to the true proportions of adaptive substitutions than are the estimates generated from count data, but accurately estimating α is notoriously difficult (Eyre-Walker, 2002; Messer & Petrov, 2013; Stoletzki & Eyre-Walker, 2011) and polyDFE may possess its own biases. In any case, it is the relative patterns between classes of genes within a species, and not the absolute value of α, that is of primary significance here.

*Information on newly sequenced datasets*

In the tables below we provide additional details for our newly generated datasets. For the new, whole-genome resequencing of *Manduca sexta*, we provide alignment rate to the reference genome, mean coverage depth, and accession numbers to the raw data (Table S1). For the *Danaus plexippus* RNA-seq data, we provide raw read counts for each tissue from each male and accession numbers (Table S2).

**Supporting Tables and Figures**

| Sample | Bowtie 2 alignment | Stampy alignment | Mean coverage depth |
| --- | --- | --- | --- |
| S32 | 94.12% | NA | 18.13 |
| S33 | 93.90% | NA | 19.82 |
| S34 | 93.84% | NA | 18.90 |
| S35 | 94.02% | NA | 16.93 |
| S36 | 94.04% | NA | 19.13 |
| S37 | 94.06% | NA | 20.08 |
| S38 | 94.08% | NA | 22.44 |
| S39 | 94.01% | NA | 26.16 |
| S40 | 93.79% | NA | 19.72 |
| S42 | 94.12% | NA | 25.67 |
| S44 | 94.07% | NA | 22.10 |
| S45 | 94.16% | NA | 20.49 |
| Q6 | 78.59% | 85.44% | 11.2 |

**Table S1.** Basic alignment summary statistics of newly sequenced *Manduca* samples. Those starting with S are from *M. sexta*, Q from *M. quinquemaculata*. Alignments with bowtie2 were carried out using the parameter defaults for very-sensitive-local alignment. In order to improve alignment of heterospecific reads, stampy was used, with default parameters excepting the substitution rate parameter, which was increased to 0.1. Coverage depth was calculated with deeptools. Raw sequences can be found on NCBI with the following accession: SRP144217.

| Sample | Head | Thorax | Midgut | Testes | Accessory gland |
| --- | --- | --- | --- | --- | --- |
| M1 | 7,038,350 | 6,339,933 | 3,999,656 | 5,958,899 | 7,021,055 |
| M2 | 6,835,543 | 5,418,564 | 7,074,743 | 10,350,175 | 7,453,113 |
| M3 | 6,471,153 | 6,469,699 | 10,497,431 | 6,935,772 | 7,984,240 |

**Table S2.** Raw read counts per tissue for each sample generated for expression analyses. Tissues were taken from a lab colony of monarch butterflies at the University of Kansas for RNA extraction and sequencing. Reads were taken through standard expression analysis pipeline with Trinity to generate FPKM values for each gene within a tissue. Tissue specificity metrics were calculated with custom R scripts. Data can be accessed with accessions SRR8580831 - SRR8580842.

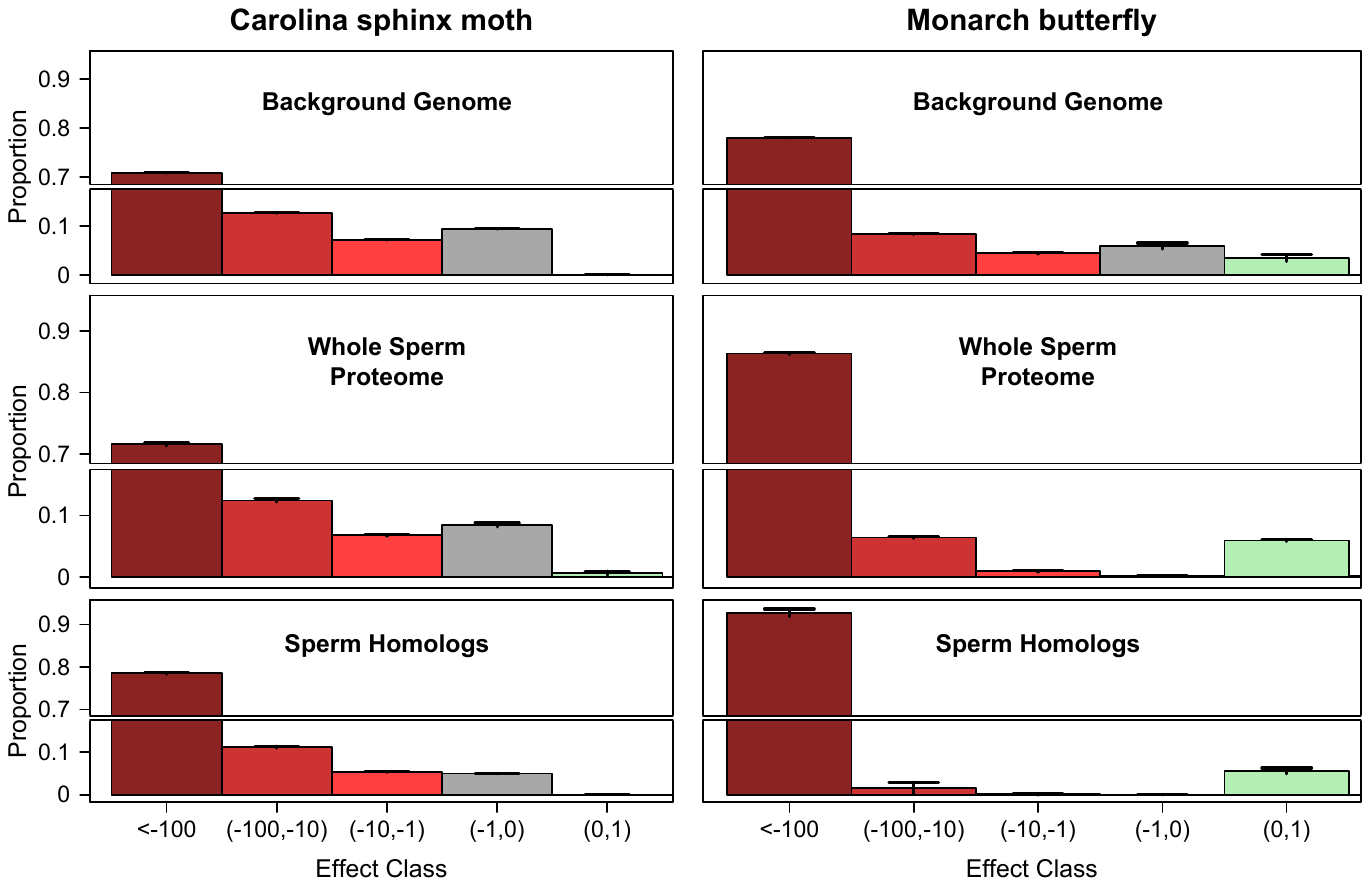

Figure S1. Predicted distribution of fitness effects of new non-synonymous mutations for the gene sets investigated in Figures 1 and 2. From top to bottom: autosomal background genome, autosomal sperm proteome, and the subset of sperm homologs found in both species. Bars represent the mean proportion for each selective class, with error bars representing twice the standard error of the mean from 100 bootstrap replicates of the input data. Note the gap in the y-axis due to the preponderance of strongly deleterious (s <-100) mutations. Left. The DFE shows little variation between the background and the sperm data sets, baring a slight increase in the proportion of strongly deleterious mutations. Right. In monarch butterflies, note the increasingly bimodal distribution of fitness effects that coincides with increased selection inferred from earlier analyses. In the sperm proteome (middle right box), there is a decrease in effectively neutral (gray, -1 < s < 0) and weakly deleterious (light red, -100 < s < -1) variants, with a concomitant increase in both strongly deleterious (dark red, s < -100) and beneficial (green, 0 < s < 1) variants. In sperm homologs this effect is even more pronounced, with nearly all variants under selection.

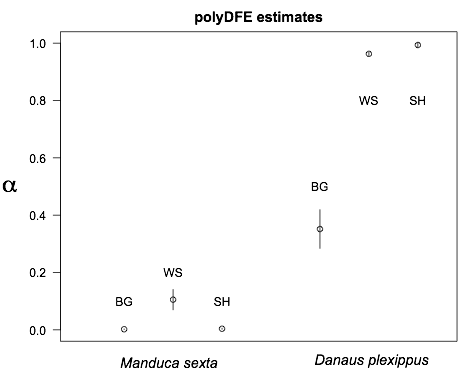

Figure S2. Estimates of α from polyDFE. Points are the mean parameter estimate from 100 bootstrap replicates of the input site frequency spectra for three classes of genes: the genome background (BG), whole sperm proteome (WS), and sperm homologs (SH) in each species. Bars represent twice the standard error of the estimate. Note that for some estimates (e.g. *Manduca* genome background, likelihood nearly always converged on the same estimate, leading to low variance). While more adaptive evolution is detected in the sperm proteome of *Manduca* than the genome background, the magnitude of the difference is dwarfed by the difference in monarchs.

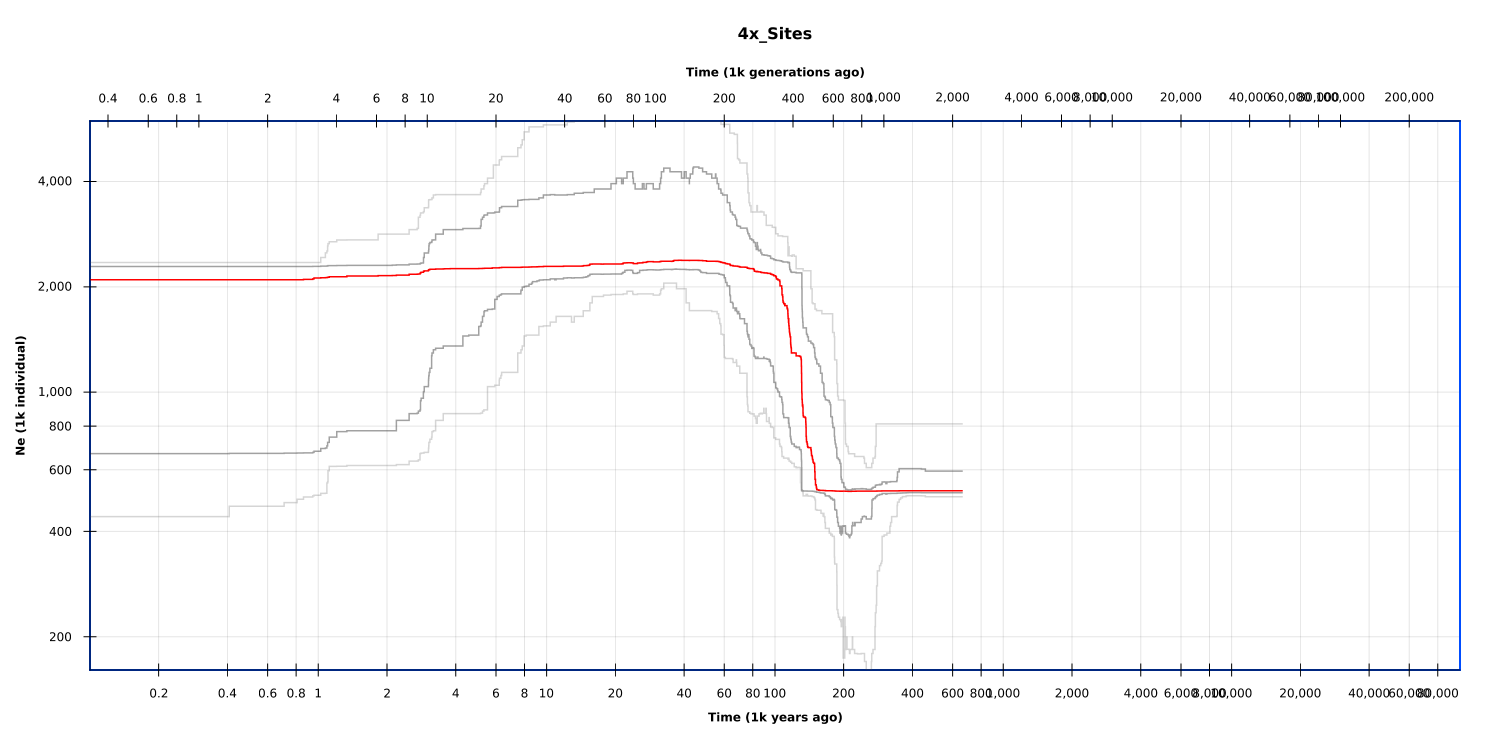

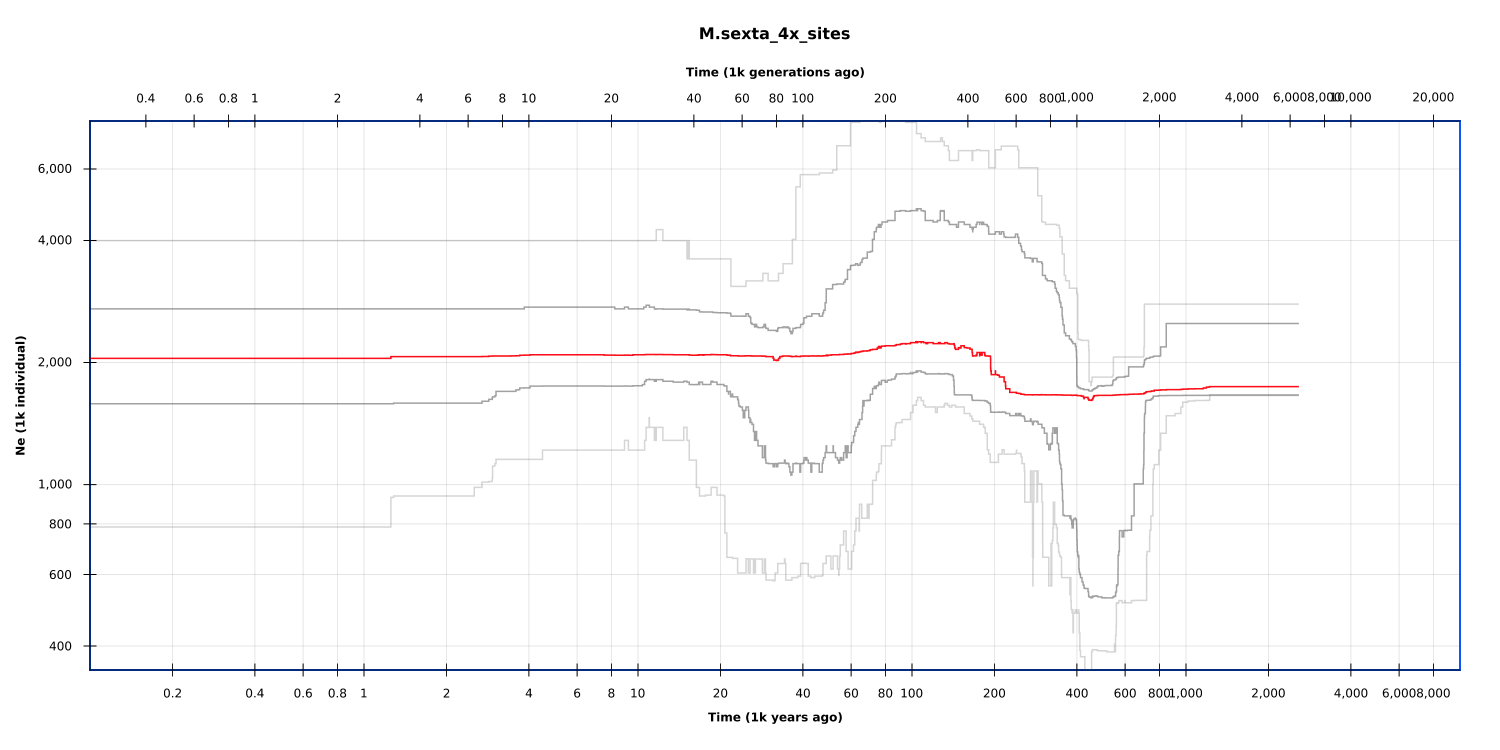

Figure S3. Inferred population sizes and histories for both monarchs (top) and Carolina sphinx moths (bottom) based on coalescent history of 4-fold degenerate sites with the program stairway plot using the estimated mutation rate from *Heliconius melpomene*. Red lines represent mean of estimates, dark and light grey bars represent 90% and 95% confidence intervals. +
